# Global deletion of *Malat1* alters alcohol consumption in a sex-specific manner

**DOI:** 10.64898/2026.06.19.733448

**Authors:** Daniela V. Gil, Annalisa M. Baratta, Carolyn Ferguson, Martha Miskanic, Andrew Iker, Gregg E. Homanics, Sean P. Farris

**Affiliations:** Center for Neuroscience at the University of Pittsburgh, Pittsburgh, PA, USA; Department of Anesthesiology & Perioperative Medicine, University of Pittsburgh, Pittsburgh, PA, USA; Department of Neurobiology, University of Pittsburgh, Pittsburgh, PA, USA; Department of Pharmacology & Chemical Biology, University of Pittsburgh, Pittsburgh, PA, USA; Department of Biomedical Informatics, University of Pittsburgh, Pittsburgh, PA, USA

## Abstract

Alcohol use disorder (AUD) is a widespread psychiatric condition, yet the molecular mechanisms underlying its development remain poorly understood. While prior studies have largely focused on protein-coding genes, long non-coding RNAs (lncRNAs) remain underexplored in AUD. *Malat1*, a highly abundant and evolutionarily conserved lncRNA, is elevated in post-mortem brain tissue of human AUD subjects and rodents chronically exposed to ethanol; however, its causal contribution to AUD-relevant behaviors remains unknown. Using CRISPR/Cas9 genome editing, we generated two complementary global *Malat1* knockout models to assess its role in alcohol intake and related phenotypes. Constitutive knockout selectively attenuated acute functional tolerance rate and every-other-day two-bottle-choice alcohol intake in females. These results were supported by an inducible adult conditional global knockout model, which reduced ethanol consumption in females without altering taste preference. Together, our findings provide the first causal evidence that *Malat1* regulates alcohol consumption in a sex-specific manner, supporting further investigation into its underlying mechanisms in AUD.

## 1. Introduction

Alcohol use disorder (AUD) is a psychiatric disorder characterized by an inability to stop or control alcohol use despite adverse social, occupational, or health consequences.^1^ Approximately 10% of the US population aged 12 and older meet the criteria for AUD,^2^ and chronic excessive alcohol use can be attributed to over 117,000 annual deaths,^3^ underscoring the need to identify the neuroadaptations underlying AUD.

Although most studies focus on highlighting the roles of proteins in AUD pathology, protein-coding genes comprise only 2% of the human genome,^4^ leaving much of the genome underexplored. Recent studies have identified long non-coding RNAs (lncRNAs), as regulators of biological processes underlying various diseases such as cancer, neurodegenerative disorders, and substance use disorders.^5–12^ LncRNAs are transcripts longer than 200 nucleotides that do not encode proteins, but can interact with DNA, RNA, and proteins to regulate cellular and molecular functions.^13–15^ LncRNA genes outnumber protein-coding genes in human and mouse genomes (approximately 36,000 versus 20,000);^16^ nevertheless, lncRNAs remain challenging to study due to limited evolutionary conservation and high tissue- and cell-type specificity.^17–19^

*Metastasis associated lung adenocarcinoma* (*Malat1*), also known as *Neat2* (*nuclear enriched abundant transcript 2*), is one of the most evolutionary conserved and abundant lncRNAs.^17,20–22^ Among its numerous biological functions, *Malat1* interacts with transcription factors to regulate synapse formation,^21,23,24^ gene expression and alternative splicing,^13,25–27^ and also modulates proinflammatory cytokine release and cell death pathways.^7,28–30^

Growing evidence suggests chronic alcohol exposure triggers immune responses in the peripheral and central nervous system (CNS), which may, in turn, promote alcohol consumption and promote dependence in a framework called the neuroimmune hypothesis of AUD.^31–35^ Although a causal role for *Malat1* in AUD has not been established, *Malat1* modulates several inflammatory pathways linked to alcohol-induced inflammation, alcohol consumption and other alcohol-related behaviors and pathologies.^36–60^ Among these, *Malat1* positively regulates the Hmgb1/Tlr4/NF-κB axis,^61–64^ Nlrp3,^65–68^ and Pde4 activity, ^68–73^ directly or through microRNA (miRNA) repression. Through these pathways, *Malat1* may represent an upstream coordinator of the neuroimmune signaling underlying in AUD pathogenesis. Notably, *Malat1* was found elevated in post-mortem brains of individuals with AUD^74^ and in rodents chronically exposed to alcohol.^74–76^

Given *Malat1*’s role in promoting neuroinflammatory signaling, we hypothesized that *Malat1* may regulate alcohol consumption through modulation of neuroimmune activity, and predicted *Malat1* knockout would attenuate alcohol intake. To test this, we used the CRISPR Turbo Accelerated Knockout (CRISPy TAKO)^77^ approach to generate a cohort of *Malat1* knockout mice to rapidly screen for *Malat1*-mediated effects on alcohol-related behaviors. To address potential confounds of developmental compensation in the CRISPy TAKO mice, we also developed a Cre-dependent, conditional *Malat1* global knockout (cKO) line, induced with tamoxifen in adulthood. Using a behavioral test battery, we observed female-specific reductions in acute functional tolerance rate and non-dependent alcohol consumption in *Malat1* TAKOs using an every-other-day two-bottle-choice (EOD-2BC) paradigm, which were partially replicated in the inducible *Malat1* global cKO model.

## 2. Materials and Methods

### 2.1 Animals

All experiments were approved by the Institutional Animal Care and Use Committee (IACUC) of the University of Pittsburgh and conducted in accordance with the National Institutes of Health Guidelines for the Care and Use of Laboratory Animals. All animals were housed in facilities accredited by the American Association for the Accreditation of Laboratory Animal Care International. Mice were maintained on a 12/12-hour light-dark cycle (lights on at 0700) and provided *ad libitum* access to food and water.

Adult, male and female C57BL/6J (B6J) mice (Strain# 000664) were purchased from The Jackson Laboratory (Bar Harbor, ME) for embryo generation and controls. Adult CD-1 recipient females and vasectomized males (Strain# 022) were obtained from Charles River Laboratories Inc. (Wilmington, MA).

### 2.2 Generation of Malat1 Turbo Accelerated Knock Out (TAKO) Mice

#### gRNA Design to Target *Malat1* Promoter

*Malat1* TAKO mice were developed using the CRISPy TAKO method, as described previously.^77^ Four guide RNA (gRNA) target sequences flanking the *Malat1* promoter^78^ (Supplementary Table 1) were identified using CRISPOR software.^79^ Custom CRISPR RNAs (crRNAs) were generated using the ALT-R™ CRISPR/Cas9 Genome Editing System (IDT DNA, Coralville, IA), and individually annealed with an invariant trans-activating crRNA (tracrRNA) at a 1:2 molar ratio to form functional gRNAs.^80^ These four gRNAs were then combined into a single solution to increase mutagenesis efficiency.^77^ Embryo collection, electroporation, and implantation were performed as described previously.^77^

### 2.3 Generation of Malat1 Floxed Animals and Breeding to Cre Mice

To create a *Malat1* floxed mouse line on a B6J background, CRISPOR was first used to design a gRNA targeting ∼3kb upstream of the transcriptional start site (*Malat1* 5’ sgRNA, Supplementary Table 1). A single-stranded oligonucleotide vector containing a loxP site, a BamH1 restriction site, and 3’ and 5’ arms each with 50 base pairs of homology were designed to target the same location (**Figure 3A**). Embryo collection, electroporation, and implantation were performed as described previously.^77^ Offspring were analyzed for precise insertion of the 5’ loxP site by PCR and Sanger sequencing (**Figure 3B**). Once offspring containing the 5’ loxP site were identified and reached adulthood, three male mice containing the 5’ loxP site were bred to wild-type (WT) B6J females, embryos were collected, and a 3’ loxP site inserted using a *Malat1* 3’ gRNA (Supplementary Table 1) and loxP insertion oligo that specifically target the *Malat1* gene ∼1kb downstream of the polyA site. These embryos were inserted into pseudopregnant females and produced four potential founder animals with both 5’ and 3’ loxP sites. Precise insertion of both loxP sites was verified by Sanger sequencing and PCR (**Figure 3B,D**). Primers used to verify the presence of both loxP sites are listed in Supplementary Table 2. Two potential founders were test-mated and produced offspring with loxP sites on the same chromosome (*i.e.,* in cis). A single founder animal with 5’ and 3’ loxP sites in cis was chosen to establish the colony by mating to WT B6J mice. Possible off-target sites for the 5’ and 3’ loxP insertions were predicted using CRISPOR, and the top 14 and 15 candidates were amplified from founder DNA. All predicted off-target sites were wildtype except one, which failed to amplify and was not pursued further. Heterozygous floxed offspring were mated to produce *Malat1* homozygous floxed mice.

Homozygous floxed mice were bred to UBC-CreERT2 mice (Strain# 007001, The Jackson Laboratory) to produce a colony of tamoxifen-inducible, *Malat1* global cKO animals. For experimental studies, breeding pairs consisted of homozygous floxed, hemizygous UBC-CreERT2 animals mated to homozygous floxed, Cre-negative animals. Sex of Cre-positive breeders were balanced across pairs to minimize potential sex-linked confounds. The presence of Cre was verified by Transnetyx (Cordova, TN).

### 2.4 Tamoxifen-Induced Recombination

Five- to seven-week-old male and female homozygous floxed, hemizygous UBC-CreERT2 mice received 150 mg/kg tamoxifen (Sigma-Aldrich, #T5648-5G; dissolved in 20 mg/mL in sterile corn oil) or vehicle (Sigma-Aldrich, C8257-500ML) i.p. once daily for five consecutive days. Mice were allowed to recover undisturbed for 10 days before starting behavioral testing. Body weight was monitored daily during tamoxifen treatment, and weekly following recovery. Validation of recombination following tamoxifen treatment was performed by RT-qPCR of RNA extracted from cortical and cerebellar tissue (see below).

### 2.5 Genotyping

To validate embryo mutagenesis in TAKO mice, DNA was collected and amplified from individual embryos using the Repli-G kit (Qiagen, #150025). To genotype TAKO and floxed offspring, DNA was extracted from ear clips using Quick Extract (Lucigen, #QE09050). Ear clips from cKO mice were also collected at sacrifice to verify peripheral recombination. DNA samples were subjected to PCR using the following protocol: 95°C for 5 min (1x); 95°C for 30 s, 60°C for 30 s, 72°C for 1 min (40x); 72°C for 10 min (1x). Amplicons were analyzed by agarose gel electrophoresis and Sanger Sequencing. PCR primers are listed in Supplementary Table 2.

### 2.6 Behavioral Assays

Upon reaching six weeks of age, *Malat1* TAKO mice underwent a battery of ethanol-related behavioral tests to examine the impact of *Malat1* deletion on behavior. All TAKO animals underwent drinking-in-the-dark (DID) followed by every-other-day, two-bottle-choice (EOD-2BC) drinking. A subset then performed the acute functional tolerance test (AFT) while a different group underwent ethanol clearance testing. A one-week washout period occurred between each test.

Seven- to nine-week-old *Malat1* cKO mice underwent EOD-2BC drinking upon recovery from tamoxifen-induced recombination.

#### Drinking-in-the-Dark (DID)

Animals were single-housed and received one sipper tube made from a 10mL serological pipette containing 20% (v/v) ethanol (Decon Labs, Inc., King of Prussia, PA, USA) beginning 3h into the dark cycle. Following 2h of access, ethanol volume consumed was measured, the tube removed, and animals given access to water. Water was again removed 24h after starting the first test, and the test repeated with 4h ethanol access, after which ethanol volume consumed was measured and a blood sample was immediately collected from the tail vein for assessment of blood ethanol concentration (BECs). Ethanol consumption was calculated for both days (grams ethanol consumed/kg body weight). Ethanol concentration in serum was assessed using an Analox Alcohol Analyzer (AM1, Analox Instruments, London, UK).

#### Acute Functional Tolerance (AFT)

Mice were trained to remain on a stationary 1-inch diameter rotarod (Ugo Basile, Province of Varese, Italy) for one minute prior to beginning the test. Animals received 1.75 g/kg i.p. ethanol, were placed back on the rotarod, and latency to fall was recorded. Animals were placed back on the rotarod every few minutes until they could successfully balance for 30 seconds. Time to regain balance was recorded (T1) and blood samples for BEC analysis were collected from the tail vein (BEC1). A second injection of 2.0 g/kg i.p. ethanol was given and mice were tested until they could remain on the rotarod for 30 seconds. The time was recorded (T2), and a second blood sample was collected from the tail vein (BEC2). Acute functional tolerance (AFT) was determined as BEC2 – BEC1, and AFT rate was calculated as BEC2 – BEC1/T2 – T1.

#### Ethanol Clearance

Animals were i.p. injected with 3.5 g/kg ethanol. Blood was collected from the tail vein 30, 60, 120, and 180 minutes after injection for assessment of BECs.

#### Every-Other-Day, Two-Bottle-Choice (EOD-2BC) Drinking

Mice were single-housed three days prior to drinking, and given two water bottles (Amuza, #H37.100.03) equipped with ball-bearing sippers (Amuza, #H37.100.0) to acclimate to experimental conditions. Access to rack-mounted Lixits was removed during this period.

Upon starting EOD-2BC, mice received alternating 24h access to either water and ethanol, or water only. Ethanol was presented in escalating concentrations (3%, 6%, 9%, and 12% for 1 day each; 15% for 3 days; 20% for 8 days, v/v), with ethanol bottle position switched each session to prevent side preferences. Bottle weights were recorded every 24h, and total fluid intake, ethanol intake, and ethanol preference were calculated after each session. To control for evaporation and drippage, identical bottles were placed in two empty cages and weighed alongside experimental cages. Daily intake values were excluded if technical errors occurred (*e.g.,* bottle leakage, Lixit devices not removed).

After the final EOD-2BC session, *Malat1* cKO and control mice underwent a seven-day washout period followed by continuous 2BC access to saccharin (Sigma-Aldrich, #240931-50G) to determine changes in sweet taste preference. Saccharin concentration increased over 8 days (2 days each at 0.0066%, 0.0165%, 0.033%, and 0.066%). Following another seven-day washout period, the experiment was repeated with quinine (Sigma-Aldrich, #145912-10G), a bitter tastant (2 days each at 15 µM, 30 µM, and 60 µM).

### 2.6 Tissue Collection

Mice were euthanized with CO_2_, and whole brains fresh-frozen in liquid nitrogen. Samples were maintained at −80°C until processing for RNA extractions.

### 2.7 RNA Isolation and RT-qPCR

Cortical and cerebellar tissue were used for RT-qPCR analysis. Total RNA was isolated using Trizol (Invitrogen, #15596018) according to manufacturer’s instructions. RNA was purified using the TURBO DNA-free™ Kit (Invitrogen, #AM1907) and purity and concentration were assessed with a Nanodrop Spectrophotometer (Thermo Scientific, Waltham, MA). RNA (500 ng) was converted to cDNA with the Superscript™ III First-Strand Synthesis System (Invitrogen, #18080051) and used for RT-qPCR with iScript SYBR green (BioRad, Hercules, CA) on a BioRad iCycler. Expression was calculated with the 2^−ΔΔCt^ method^81^ and normalized to β-actin. RT-qPCR primers are listed in Supplementary Table 2.

### 2.8 Statistics

Statistical analyses were conducted in GraphPad Prism (La Jolla, CA). Normality was first assessed using the Shapiro-Wilk test, and outliers were identified using the ROUT test (Q = 1%). Data with normal distribution were analyzed using one-tailed, unpaired student’s t-tests or Welch’s t-tests (genotype comparisons), or parametric 2-way ANOVAs or mixed-effect models (genotype x time [repeated measure]). Data not normally distributed were analyzed using the non-parametric Mann-Whitney test.

## 3. Results

### 3.1 Generation of *Malat1* CRISPy TAKO Mice

Four gRNAs were used to target *Malat1*, with two upstream and two downstream of the transcriptional start sites (**Figure 1A)**. F0 CRISPy TAKO mice derived from electroporated embryos were genotyped by PCR to assess CRISPR-induced mutations (**Figure 1B**). The wild-type (WT) allele produced a 933 base pair amplicon, whereas edited alleles exhibited smaller bands consistent with deletions in the targeted region. PCR results revealed variable deletion patterns across animals, suggesting heterogeneous CRISPR-mediated gene editing at the targeted locus as expected. Mice exhibiting WT bands were excluded from further experimentation.

**Figure 1.**
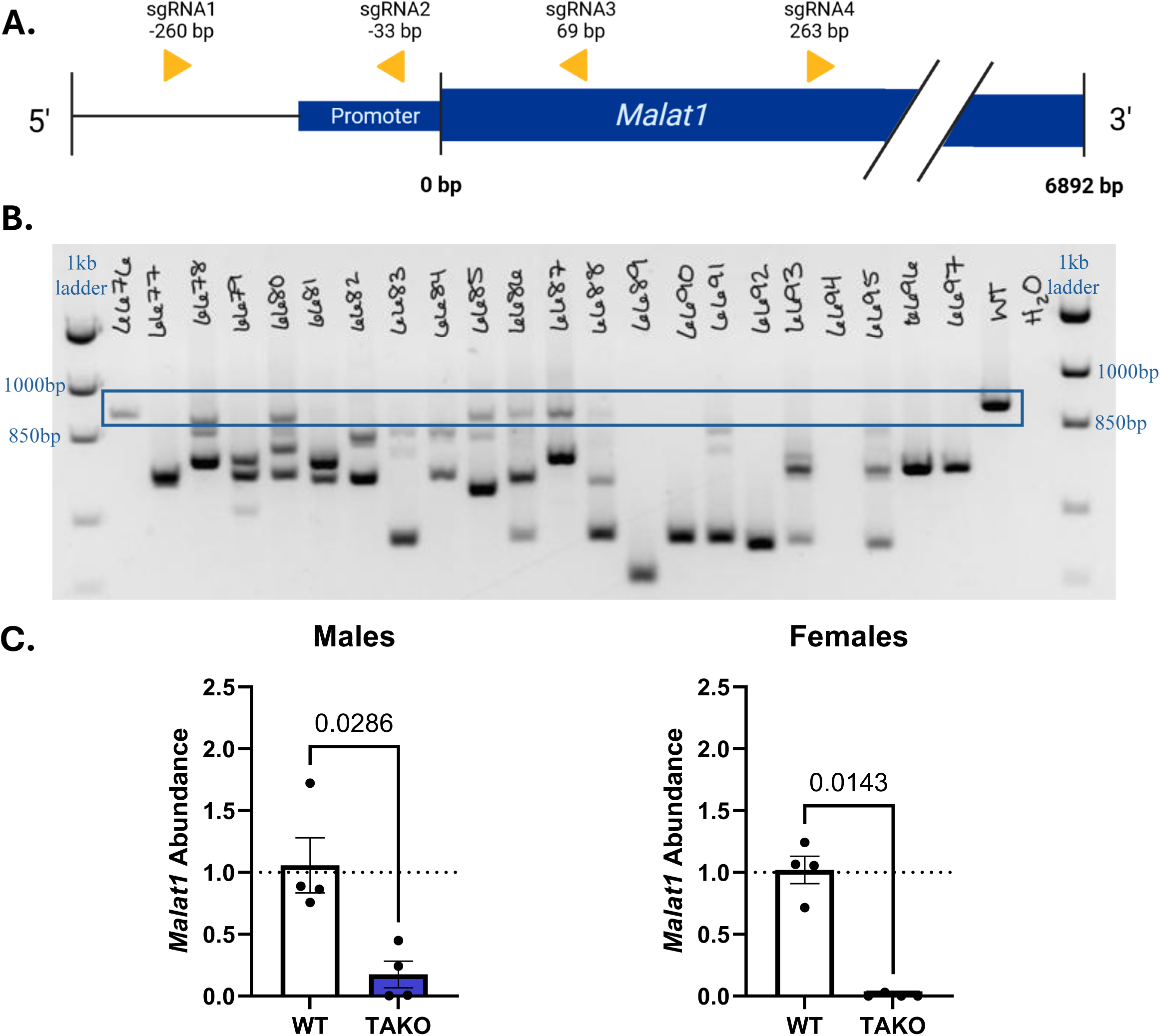
Generation of *Malat1* CRISPy TAKO mice. **(A)** gRNA target sites surrounding *Malat1* promoter. **(B)** PCR results from DNA of *Malat1* CRISPy TAKO mice. Animals showing WT bands (boxed; 933 bp amplicon) were excluded from behavioral testing. **(C)** RT-qPCR results from cortical tissue showing reduced relative abundance of *Malat1*, normalized to *β-actin*. Values represent mean ± SEM. n= 4/group.

To validate functional *Malat1* knockout, cortical tissue was collected from a cohort of adult offspring and analyzed by RT-qPCR. Male and female *Malat1* CRISPy TAKO mice exhibited significantly reduced *Malat1* abundance compared to WT controls (males: U= 0, p= 0.0268; females: U= 0, p= 0.0143, Mann-Whitney), with transcript abundance reduced to approximately 0-20% of WT levels (**Figure 1C**).

Together, these findings demonstrate that CRISPy TAKO-mediated targeting robustly reduced *Malat1* abundance, confirming the generation of functional *Malat1* TAKO mice for behavioral screening.

### 3.2 Behavioral Assessment of TAKOs

To determine *Malat1*’s effects on ethanol-related phenotypes, *Malat1* TAKO mice underwent a battery of behavioral testing, assessing binge-drinking, ethanol clearance, acute functional tolerance, and non-dependent voluntary drinking.

A two-day DID paradigm was used to assess binge-like ethanol consumption. Mice were given access to a sipper bottle containing 20% ethanol (v/v) three hours into the dark cycle. Female *Malat1* TAKOs drank significantly more ethanol after two-hour access on day one (t= 1.870, p= 0.0370, Student’s t-test), whereas males exhibited no changes in drinking (t= 1.561, p= 0.0657, **Figure 2A**). However, intake was unaltered in both sexes following four-hour access on day two of DID (males: t= 0.8579, p= 0.1990, Welch’s t-test; females: t= 1.149, p= 0.1300; Student’s t-test, **Figure 2A,B**). Blood collected immediately following four-hour exposure showed similar BECs in *Malat1* TAKO males (U= 74.50, p= 0.0733, two-tailed Mann-Whitney) and females (t= 0.5174, p= 0.6088, two-tailed Student’s t-test; **Figure 2A,B**) compared to WT controls.

**Figure 2.**
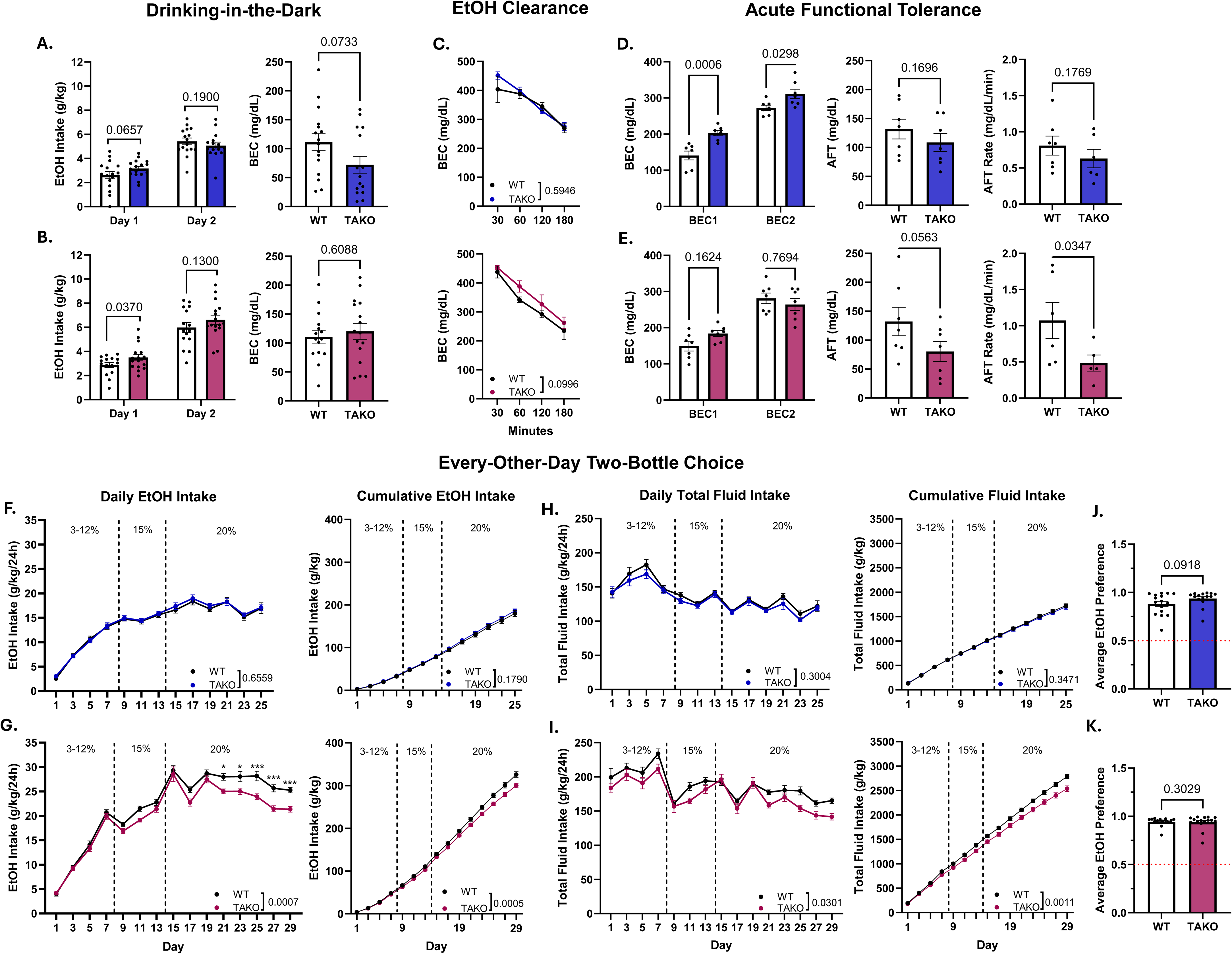
*Malat1* CRISPy TAKO behavioral battery. **(A, B)** Drinking-in-the-Dark (DID) ethanol intake (left) and blood ethanol concentrations (BECs; right) in males (A) and females (B). Mice were given access to ethanol for two hours on day one, and four hours on day two. No changes in ethanol intake or BECs were observed in either sex on day two. **(C)** No effect of genotype in either sex was observed on ethanol clearance from blood following i.p. injection with 3.5 g ethanol per kg body weight. **(D,E)** Acute functional tolerance (AFT) assay in males (D) and females (E). BECs recorded upon first (BEC1) and second (BEC2) recovery of balance following ethanol injections are shown (left). Only males reported higher BEC values at both timepoints. AFT (center), calculated as BEC2-BEC1, did not differ between genotypes in either sex. AFT rate (right), calculated as AFT/(T2-T1), was reduced in females. **(F,G)** Every-other-day two-bottle choice (EOD-2BC) ethanol intake in males (F) and females (G). Daily intake across concentrations (left) and cumulative intake (right) are shown. Statistical comparisons for cumulative intake reflect the final timepoint only. Ethanol intake was reduced in females but not males. **(H,I)** Total fluid intake across the EOD-2BC paradigm in males (H) and females (I). Reductions in daily and cumulative total fluid intake were observed only in females. Statistical comparisons for cumulative intake reflect the final timepoint only. **(J,K)** Average daily ethanol preference in males (J) and females (K) did not differ between genotypes. Values represent mean ± SEM. n = 5-16/group.

To probe the impact of *Malat1* on ethanol clearance, blood was collected at 30, 60,120, and 180 minutes following i.p administration of 3.5 g/kg ethanol. A main effect of time was observed in both sexes (males: F[1.335, 9.343] = 25.72, p = 0.0003; females: F[2.226, 17.81] = 35.04, p< 0.0001; 2-way ANOVA), but no effect of genotype (males: F[1, 7] = 0.3107, p= 0.5946; females: F[1, 8] = 3.468 p= 0.0996; 2-way ANOVA), nor time x genotype interaction (males: F[1.335, 9.343] = 0.9696, p = 0.3778; females: F[2.226, 17.81] = 35.04, p= 0.2107; 2-way ANOVA, **Figure 2C**), was found, indicating *Malat1* does not alter ethanol clearance.

AFT was chosen to determine if *Malat1* influences physiological tolerance and motor coordination in response to ethanol. Following i.p injection of 1.75 g/kg, animals were placed on a rotarod and re-tested until they could balance for 30 seconds; blood was collected immediately (BEC1). A second injection of 2.0 g/kg ethanol was given, and the test was repeated and blood collected upon regaining balance (BEC2). BECs were significantly elevated in *Malat1* TAKO males at both collection timepoints (main effect of genotype: F[1, 12] = 31.34, p= 0.0001, 2-way ANOVA), though AFT and AFT rate were unaltered (AFT: t= 0.9955, p= 0.1696; AFT rate: t= 0.9680, p= 0.1769; Student’s t-test; **Figure 2D**). No genotype effect was observed on BEC in female *Malat1* TAKOs (F[1, 12] = 0.5645, p= 0.4669, 2-way ANOVA). AFT showed a trend toward reduction (t= 1.728, p= 0.0563; Student’s t-test), while AFT rate was significantly reduced in female TAKOs relative to controls (t= 2.001, p= 0.0347; Welch’s t-test; **Figure 2E**). Together, these data suggest *Malat1* modulates AFT rate in females.

Non-dependent voluntary ethanol consumption was assessed using EOD-2BC. Single-housed mice were given alternating access to escalating concentrations of ethanol (3-20%, v/v) for a total of 25 (males) or 29 (females) days. Male *Malat1* TAKOs exhibited no differences in ethanol intake, either in daily measures (F[1, 30] = 0.2026, p= 0.6559, 2-way mixed-effects model) or cumulative intake at the final timepoint (U= 96, p= 0.1790, Mann-Whitney; **Figure 2F**). Total fluid intake was also unaffected at the daily (F[1, 30] = 1.110, p= 0.3004, 2-way mixed-effects model) or cumulative (t= 0.3978, p= 0.3471, Student’s t-test; **Figure 2H**) level, and average ethanol preference did not differ (U= 92, p= 0.0918, Mann-Whitney; **Figure 2J**).

In contrast, female *Malat1* TAKOs exhibited reduced ethanol consumption at both the daily (F[1, 29] = 14.43, p< 0.0001, 2-way mixed-effects model) and cumulative (t= 3.657, p= 0.0005; Student’s t-test; **Figure 2G**) levels. Post hoc analysis of the significant genotype x time interaction (F[14, 405] = 2.962, p= 0.0002, 2-way mixed-effects model) revealed that *Malat1* TAKOs consumed significantly less ethanol than controls in the final five days of 20% ethanol access (all p< 0.05; Bonferroni’s). This effect, however, was accompanied by reduced total fluid intake both daily (F[1, 29] = 5.204, p= 0.0301, 2-way mixed-effects model) and cumulatively (t= 3.433, p= 0.0011; Welch’s t-test), with no observed differences in average ethanol preference (t= 0.2152, p= 0.4156; Student’s t-test). Together, this suggests female-specific effect of *Malat1* cKO on ethanol-related behaviors.

### 3.3 Generation of Conditional *Malat1* Global Knockouts

To ensure *Malat1*-mediated changes in ethanol-related behaviors were not driven by compensatory developmental effects, we used Cre/loxP technology to create an inducible global knockout model. Briefly, CRISPR/Cas9 genome editing technology was used to insert loxP sites upstream of the transcriptional start site and downstream of the polyadenylation site (**Figure 3A**), and insertion was verified with Sanger sequencing of PCR amplicons (**Figure 3B**). Floxed animals were bred to a UBC-CreERT2 line to ultimately create tamoxifen-inducible homozygous *Malat1* floxed, hemizygous UBC-CreERT2 *Malat1* cKOs.

**Figure 3.**
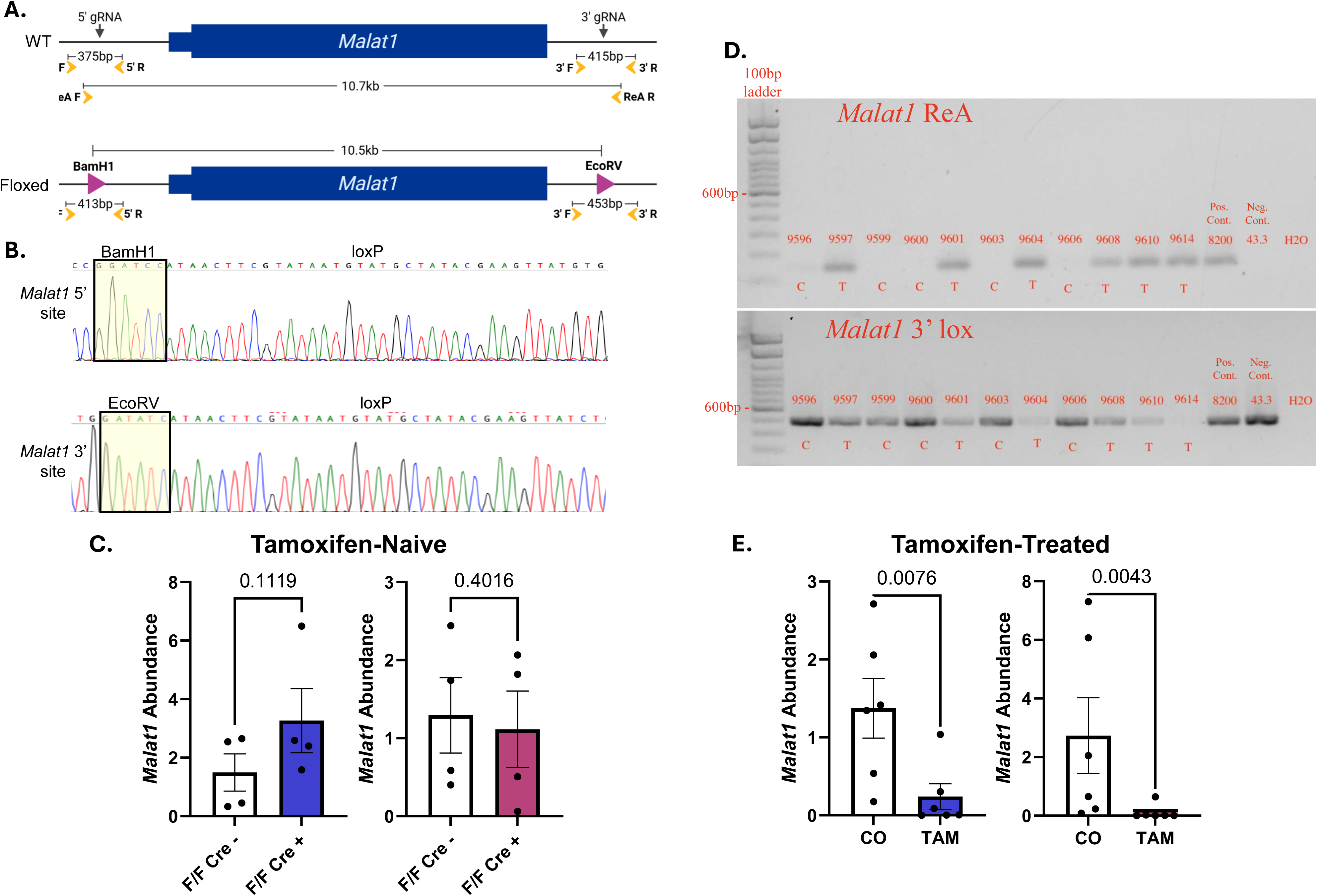
Generation of *Malat1* cKOs. **(A)** Schematic representation of the WT and *Malat1* floxed alleles. loxP sites (pink) flank the *Malat1* locus. Yellow arrows represent PCR primers. **(B)** Sanger sequencing chromatograms demonstrating 5’ and 3’ loxP site insertion into the DNA. **(C)** RT-qPCR results showing no changes in cortical *Malat1* abundance between tamoxifen-naive homozygous-floxed *Malat1* mice lacking Cre (F/F Cre-) and those expressing hemizygous UBC-Cre (F/F Cre+). RT-qPCR results are normalized to *β-actin*. **(D)** Validation of UBC-CreERT2-mediated recombination in ear clips of corn oil- (C) or tamoxifen-treated (T) mice. PCR with recombination-specific (ReA) primers detect Cre-mediated excision and *Malat1* 3’ lox primers detect the floxed allele. Samples with known recombination statuses were used as positive and negative controls, respectively. **(E)** RT-qPCR results showing reduced cortical *Malat1* abundance in F/F Cre+ mice following treatment with tamoxifen compared to corn oil (CO). RT-qPCR results are normalized to *β-actin*. Values represent mean ± SEM. n= 4-6/group.

Preliminary analyses verified loxP sites did not alter *Malat1* levels compared to wildtype B6J mice (data not shown). To assess whether Cre altered baseline (*i.e.*, prior to tamoxifen-induced recombination) *Malat1* levels, RT-qPCR compared cortical *Malat1* abundance in tamoxifen-naïve homozygous floxed mice, with or without Cre (Cre+ and Cre-, respectively). No differences in *Malat1* abundance were observed between Cre- and Cre+ mice (males: t=1.396, p= 0.1119, Welch’s t-test; females: t= 0.2604, p= 0.4016, Student’s t-test; **Figure 3C**), indicating loxP insertion and Cre expression do not impact *Malat1* levels independently of tamoxifen.

Next, functional recombination of the *Malat1* locus was validated using PCR and RT-qPCR. PCR analysis of genomic DNA from ear tissue detected recombination only in tamoxifen treated animals (**Figure 3D**) demonstrating tamoxifen-induced deletion of the Malat1 gene. In contrast, no recombination was detected in samples from corn oil treated mice. However, the intact unrecombined floxed allele was detected in both corn oil- and tamoxifen-treated Cre+ mice (**Figure 3D**), demonstrating that tamoxifen-induced recombination was incomplete in the tamoxifen treated animals. Despite the presence of a low level of unrecombined Malat1 following tamoxifen treatment, RT-qPCR analysis of cortical tissue revealed a significant reduction in *Malat1* abundance in tamoxifen-treated homozygous floxed, Cre+ mice compared to corn-oil controls, confirming tamoxifen-dependent functional disruption of the *Malat1* gene (males: U= 3, p= 0.0076; females: U= 2, p= 0.0043; Mann-Whitney; **Figure 3E**).

### 3.4 *Malat1* cKO EOD-2BC Drinking

EOD-2BC was assessed in *Malat1* cKO mice to determine if the female-specific reduction in ethanol intake observed in TAKOs would be recapitulated in an inducible model. Ten days after treatment with corn oil or tamoxifen, ethanol-naive cKO mice underwent EOD-2BC as described previously.

Male cKOs exhibited no differences in ethanol consumption in daily measures (F[1, 30] = 0.05267, p= 0.8200, 2-way mixed effects model) or at the cumulative timepoint (t= 0.3795, p= 0.3541, Welch’s t-test; **Figure 4A**), nor in average ethanol preference (U= 115, p= 0.3434, Mann-Whitney; **Figure 4E**) relative to controls. Total fluid intake, however, was significantly elevated at both the daily (F[1, 30] = 15.98, p= 0.0004, 2-way mixed model’s effect) and cumulative (t= 3.632, p=0.0007, Welch’s t-test; **Figure 4C**) levels in male cKOs compared to controls.

**Figure 4.**
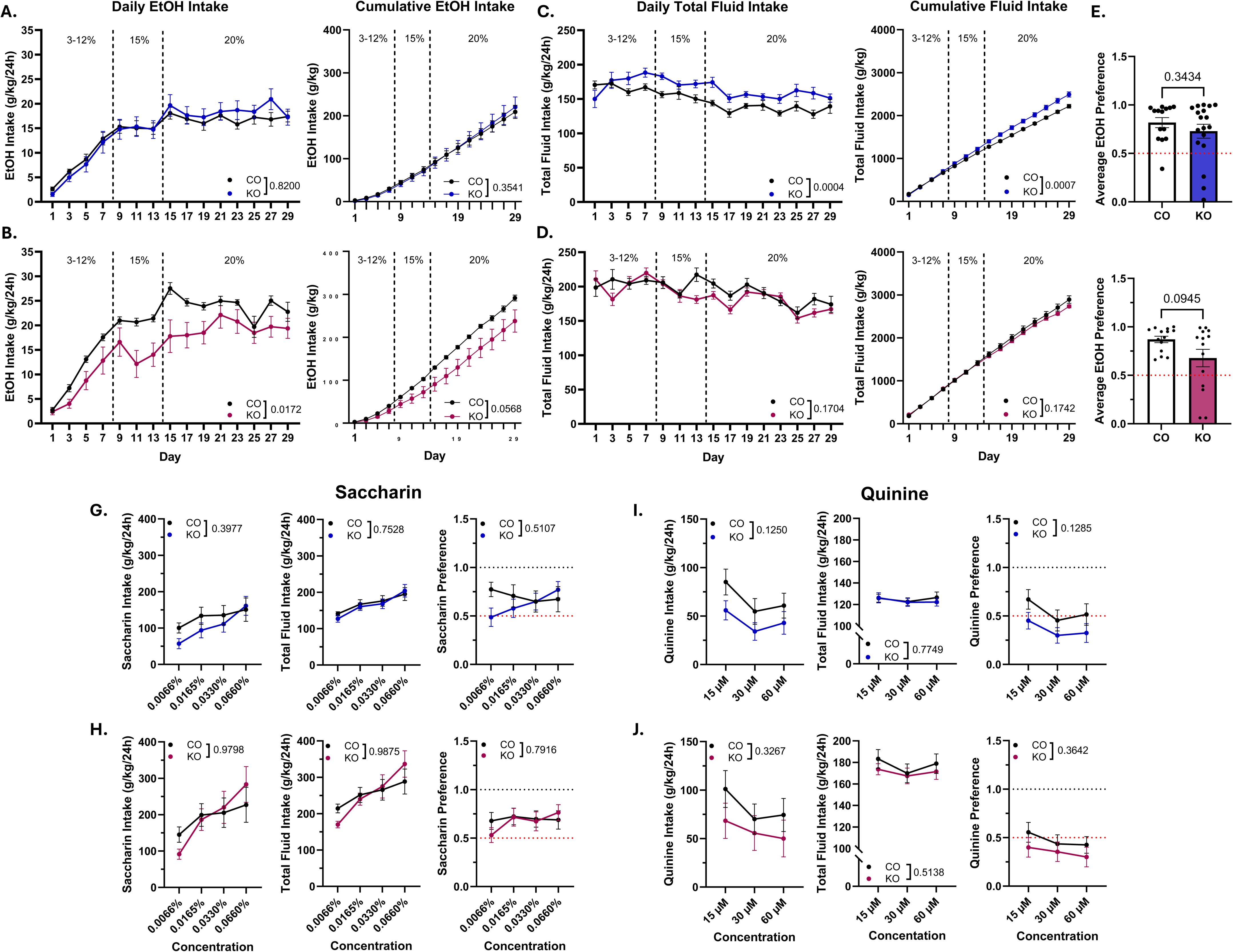
EOD-2BC drinking in *Malat1* cKO mice. **(A,B)** EOD-2BC ethanol intake in control (CO) or cKO males (A) and females (B). Daily intake across concentrations (left) and cumulative intake (right) are shown. Statistical comparisons for cumulative intake reflect the final timepoint only. Ethanol intake was reduced in females but not males. **(C,D)** Total fluid intake across the EOD-2BC paradigm in males (C) and females (D). *Malat1* cKO males increased daily and cumulative total fluid intake, whereas no differences were observed in females. Statistical comparisons for cumulative intake reflect the final timepoint only. **(E,F)** Average daily ethanol preference in males (E) and females (F). Ethanol preference was unaltered in either sex. **(G,H)** Saccharin intake (left), total fluid intake (center), and saccharin preference (right) in males (G) and females (H). No genotype differences were observed in any measure or sex. **(I,J)** Quinine intake (left), total fluid intake (center), and quinine preference (right) in males (I) and females (J). No genotype differences were observed in any measure or sex. Values represent mean ± SEM. n = 12-16/group.

In contrast, female *Malat1* cKOs displayed reduced daily ethanol intake (F[1, 26] = 6.476, p= 0.0172, 2-way mixed effects model; **Figure 4B**). Cumulative ethanol intake showed a trend toward significance (U= 44, p= 0.0568, Mann-Whitney; **Figure 4B**), likely because cumulative values were calculated only from animals with complete daily intake datasets (*i.e*., no exclusion of days with technical issues). Unlike results observed in female *Malat1* TAKOs, conditional KO in adulthood did not alter daily (F[1, 26] = 1.988, p= 0.1704, 2-way mixed effects model) or cumulative (t= 0.8626, p= 0.1742, Welch’s t-test; **Figure 4D**) total fluid intake, or average ethanol preference (U= 69, p= 0.0945, Mann-Whitney; **Figure 4F**).

To verify that observed differences in ethanol consumption were not attributed to altered taste perception, *Malat1* cKO mice were given continuous access to sweet (saccharin) then bitter (quinine) tastants following EOD-2BC. No main effects of genotype were observed in either sex across saccharin or quinine measures (all p > 0.05; **Figure 4G-J**).

Together, these data suggest *Malat1* cKO in adulthood produced sex-specific effects on ethanol drinking. Females exhibit reduced daily ethanol intake without changes in total fluid intake or taste preference, suggesting a selective effect on ethanol consumption. Males, however, showed increased total fluid intake without alterations in ethanol consumption, indicating *Malat1* may differentially regulate fluid consumption and ethanol drinking across sexes.

## 4. Discussion

The current study is the first to causally test *Malat1*’s role in regulating alcohol-related behaviors. We created two different *Malat1* KO mouse models: a CRISPY TAKO constitutive global KO model to rapidly screen for *Malat1*-mediated effects in F0 animals, and a Cre-dependent, tamoxifen-inducible *Malat1* cKO in adulthood that minimizes developmental compensation. Efficient reduction of *Malat1* abundance was observed in both models. Behavioral outcomes were modest and sex-dependent, with the greatest effects observed in EOD-2BC ethanol drinking behavior (**Table 1**). Consistently, both female TAKOs and cKOs had reduced daily and cumulative ethanol intake compared to controls, with no changes in ethanol preference. These differences in EOD-2BC ethanol consumption were not observed during the first week when low ethanol concentrations (3-12%) were tested. Genotype differences were only observed when high ethanol concentrations (15-20%) were subsequently tested. It is not clear if these altered responses are concentration or time dependent. These effects were not driven by changes in taste perception as preference to palatable and aversive tastes were unaltered in cKOs, or by changes in ethanol clearance/metabolism. In contrast, male TAKOs and cKOs showed no changes in daily or cumulative ethanol intake or ethanol preference.

**Table 1.** Summary of behavioral results relative to controls for TAKO and cKO models.

| Behavior | Model | Sex | Results |
| --- | --- | --- | --- |
| DID | TAKO | M | Ethanol intake not altered after 2h or 4h access<br>BEC not altered after 4h access |
|  |  | F | Elevated ethanol intake after 2h, but not 4h access<br>BEC not altered after 4h access |
| Ethanol Clearance | TAKO | M | No genotype effect |
|  |  | F | No genotype effect |
| AFT | TAKO | M | Elevated BECs at BEC1 and BEC2<br>AFT and AFT Rate not altered |
|  |  | F | BEC not altered at either timepoint<br>Reduced AFT and AFT Rate |
| EOD-2BC | TAKO | M | Daily and cumulative ethanol intake not altered<br>Daily and cumulative total fluid intake not altered<br>Ethanol preference not altered |
|  |  | F | Reduced daily and cumulative ethanol intake<br>Reduced daily and cumulative total fluid intake<br>Ethanol preference not altered |
|  | cKO | M | Daily and cumulative ethanol intake not altered<br>Elevated daily and cumulative total fluid intake<br>Ethanol preference not altered<br>Taste preference not altered |
|  |  | F | Reduced daily and cumulative ethanol intake<br>Daily and cumulative total fluid intake not altered<br>Ethanol preference not altered<br>Taste preference not altered |

TAKO mice were additionally tested for tolerance to the acute effects of ethanol using an AFT assay. A female specific effect was also observed on this test. TAKO females displayed reduced AFT and a reduced rate of AFT. While male TAKOs exhibited elevated BECs during AFT compared to controls, AFT and AFT rate did not differ between genotypes.

In a binge-like ethanol drinking assay (*i.e.,* DID), female TAKOs showed a small but significant increase in ethanol intake during two-hour access on day one. However, no genotype differences were observed during four-hour access on day two and BEC were similar between groups. In male TAKOs, ethanol intake and BECs were also not different from control animals in the DID assay.

While temporal control of gene expression is a strength of our study relative to prior approaches, the use of tamoxifen to induce recombination introduces limitations. Though widely used in genetic models, tamoxifen, a selective estrogen receptor modulator, may exert physiological effects independent of Cre-recombination.^82,83^ Tamoxifen is commonly administered at 75 mg/kg, however, excision of the large floxed *Malat1* locus (10.5 kb) in our cKO model required a higher dose (150 mg/kg) to produce efficient knockout. We think it unlikely that the changes observed in EOD-2BC ethanol drinking behavior in the *Malat1* cKO mice are due to the adverse effects of tamoxifen because (a) the female specific decrease in ethanol consumption observed is consistent with decreased ethanol consumption observed in tamoxifen naïve female *Malat1* CRISPy TAKO mice, (b) female cKOs treated with tamoxifen drank similarly to corn oil treated controls at low ethanol concentrations, (c) the effects of tamoxifen in female cKOs were specific to ethanol drinking as water, saccharin, and quinine drinking were unaffected, and (d) male cKOs received the same dose as females and their drinking behavior did not differ from controls.

The sex-dependent effects observed in this study reflect biological differences in *Malat1* function between males and females. *Malat1* has been linked to androgen- and estrogen-mediated pathogenesis in hormone-dependent cancers and endometriosis,^96–100^ indicating *Malat1* is both responsive to and regulated by sex hormone signaling. Despite the growing literature elucidating *Malat1’s* involvement in inflammatory pathways, little is known about sex-specific effects of *Malat1* function in non-pathological models. Biological sex is a well-established modulator of peripheral and neuroimmune responses,^101–105^ though the mechanisms underlying sexually dimorphic immune activity are complex and cannot be explained by hormones alone,^106,107^ suggesting various sex-linked factors may shape the effects of *Malat1* knockout. One study recently demonstrated *Malat1* plays a sex-specific role in Th2 cell differentiation, with *Malat1* loss impairing IL-2 receptor signaling and IL-2 mediated cytokine expression in females.^86^ *Malat1* may regulate Th2 function through Nlrp3, whose activity may be sexually dimorphic,^108–110^ via repression of miR-224-5p or miR*-*124*-*3p^32,71^ Chronic alcohol exposure is associated with a shift from Th1 to Th2 immune responses,^111–113^ and *Malat1* loss may counteract this shift by suppressing Th2-associated signaling, potentially contributing to the female-selective effects observed here. Together, these findings suggest that sex-specific regulation of *Malat1* may contribute to the female-selective effects observed here, highlighting the need for greater attention to sex-dependent mechanisms of *Malat1* function.

The mechanisms by which *Malat1* knockout impacts ethanol intake are unknown. *Malat1* interacts with DNA, RNA, and proteins to modulate various biological functions including regulation of gene expression and alternative splicing,^13,25–27^ synapse formation,^21,24^ and neuroimmune signaling;^7,28,29^ any of which may plausibly contribute to our observed phenotypes.

Ethanol’s influences on neuroimmune signaling are not exclusively driven by direct CNS action; ethanol acts systemically to drive inflammatory responses in the CNS and periphery, with many inflammatory mediators capable of crossing the blood-brain barrier to prime and sustain neuroinflammation.^42,84,85^ *Malat1* has been linked to several immune mechanisms thought to regulate ethanol-related behaviors, including the Hmgb1/Tlr4 axis, miR-375/Pde4d axis, and the *Malat1*/miR-124/Pde4b and Nlrp3 signaling cascades.^29,61,62,65,67^ Hmgb1 levels positively correlate to lifetime consumption,^42,53,56^ and elevated expression may blunt physiological responsivity to ethanol, evidenced by reduced intoxication, hypothermic response, and motor impairment following ethanol challenge; these effects are reversed with pharmacological systemic Hmgb1 inhibition.^56^ Reduced *Malat1* abundance has also been shown to lower Hmgb1 levels.^61,62,64^ The attenuated AFT rate observed in female TAKOs is consistent with this relationship, suggesting *Malat1* loss may reduce neuroimmune-driven physiological tolerance through downregulation of Hmgb1/Tlr4 signaling.

*Malat1* may also promote inflammatory signaling through repression of miR-224^29^ or miR-124,^70,71^ disinhibiting Nlrp3 and Pde4b.^29,68,70^ Nlrp3 is a regulator of inflammasome activation and has been implicated in ethanol-induced neuroinflammation and drinking,^36,46,49,50,57,86,87^ suggesting *Malat1-*dependent modulation of Nlrp3 may contribute to the behavioral changes observed here. Pde4 signaling has also been linked to ethanol-induced neuroinflammation and ethanol-related behaviors.^40,42,43^ Notably, the non-selective Pde4 inhibitor apremilast reduced ethanol intake in EOD-2BC and attenuated AFT in male and female mice,^45,60^ mirroring the reductions in ethanol consumption and tolerance observed in female TAKOs. Subtype-specific findings indicate Pde4b inhibition does not alter chronic drinking and accelerates recovery from ethanol-induced ataxia,^88^ contrasting with the attenuated AFT rate observed in female TAKOs and reduced drinking in both female models, indicating Pde4b is likely not driving these responses. Pde4d inhibition, however, transiently reduces chronic ethanol consumption and prevents acute functional tolerance, similar to our female results, though these effects were reported only in males.^88^ Our findings suggest *Malat1* may modulate ethanol sensitivity and consumption through neuroimmune signaling pathways, including Hmgb1/Tlr4, Nlrp3, and Pde4-related mechanisms, though the underlying mechanisms are yet to be tested.

*Malat1* regulates alternative splicing primarily through its interactions with serine-arginine (SR) splicing factors within nuclear speckles, where it binds to actively transcribed genes and modulates SR protein localization and phosphorylation states to regulate pre-mRNA processing and isoform production.^21,89–91^ Transcriptomic analysis of cortical tissue from male macaques chronically exposed to ethanol revealed enrichment and ethanol-responsivity of SR genes *SRSF1* and *SFSR11*, implicating SR-mediated splicing in AUD;^92^ however, as *Malat1’s* effects on *SRSF1* expression vary in direction across cell-types, ^93^ the extent to which *Malat1* regulates *SRSF1* across CNS cell-types in AUD remains unclear. Predictive modeling using genome-wide association studies data from the Collaborative Studies on Genetics of Alcoholism further implicated alternative splicing in AUD susceptibility. Notably, a skipped exon in *SRRM2*, an SR-coding gene essential for nuclear speckle integrity, ^94^ triggers nonsense-mediated decay of its protein and produces neuroinflammatory and complement system pathway enrichment,^95^ suggesting a mechanistic link between SR*-*mediated splicing and AUD.

Beyond SR proteins, proteome network analysis of *Malat1-*interacting proteins revealed enrichment of sub-clusters implicated in RNA processing and transcriptional regulation, including heterogenous nuclear ribonucleoproteins (hnRNPs).^96^ Malat1 directly binds Hnrnph1,^26,96,97^ and perturbation studies in mouse embryonic stem cells suggest a feedback relationship in which *Malat1* abundance regulates splicing factor availability and global RNA processing;^26^ however, *Malat1’s* effects on Hnrnph1 expression also vary in direction across cell-types.^93^ Hnrnph1 has been implicated in regulating alcohol-related behaviors. Notably, global *Hnrnph1* depletion attenuated the aversive properties of high-concentration alcohol and reduced intake selectively in males,^98^ contrasting with the female-specific effects observed here and indicating Hnrnph1-mediated splicing is unlikely to underlie our findings.

*Malat1* has been found to regulate synaptogenesis by recruiting SR splicing factor to nuclear speckles.^21^ Emerging evidence, however, indicates *Malat1* may also be trafficked to the synapse of cultured hippocampal and cortical neurons, where it regulates synaptic protein localization, and in one study, encodes a novel functional micropeptide involved in synapse formation, M1, through cytoplasmic translation.^21,24,99,100^ Synaptic *Malat1* may also undergo post-transcriptional modifications implicated in fear-extinction memory,^24^ suggesting nuclear and synaptic mechanisms may converge to regulate synaptic plasticity and learned behavior. Consistent with a functional role in these regions, *Malat1* abundance was elevated in hippocampal and cortical tissue of human AUD subjects and rodents chronically exposed to ethanol;^74–76^ nonetheless, the functional relevance of *Malat1*-mediated synaptic and splicing mechanisms to alcohol-related behavior remains an important avenue for future investigations.

Along with its mature transcript, the *Malat1* gene also produces a 61-nucleotide transfer RNA-like RNA called *Malat1*-*associated small cytoplasmic RNA* (*mascRNA*).^101^ *mascRNA* has shown immunoregulatory functions independently of *Malat1;*^22,102–105^ LPS exposure decreases *mascRNA* levels and upregulates *Il6* and *Tnf*, an effect reversed by *mascRNA* overexpression, suggesting *mascRNA* plays an inhibitory role in Tlr4-dependent NF-κB pathway regulation.^22,102^ Although downstream translational targets remain unclear, *mascRNA* also broadly promotes global translation, and its enrichment in peripheral immune cells suggests this effect may be relevant for immune protein synthesis.^105,106^

Despite its classification as a lncRNA, *Malat1* was recently found to encode micropeptides.^99,107^ Although no function has been identified for the unnamed 9-residue peptide,^107^ blocking translation of the 35-residue cytoplasmic peptide, M1, recapitulated the effects of *Malat1* depletion on synaptic proteins.^99^ *Malat1* may also exist as distinct isoforms with differing functional consequences; one study found deletion of the 3’ isoform region impairs retinal and corneal function in mice,^108^ while another showed *Malat1* may undergo N^6^-methyladenosine modification at the synapse, producing protein interactions that impair fear-extinction.^24^ Together, these findings underscore the complexity of *Malat1’s* biological influence, highlighting the need for targeted, mechanism-focused studies.

The *Malat1* floxed mice we created should be a valuable research tool for those interested in genetic dissection of *Malat1* function in vivo. These mice can be mated to tissue specific, cell specific, and or temporally regulated Cre expressing mice to produce a variety of conditional knockout mouse lines. Conditional KO mice such as these can be used to overcome the limitations of previously reported *Malat1* constitutive global KO mouse lines.^109,110^ This floxed allele was designed to enable Cre-mediated recombination of the *Malat1* locus to eliminate the large *Malat1* RNA, the endonucleolytic derived small *mascRNA* product,^101^ and all potential peptide products, i.e., a complete null allele.

In summary, we developed two distinct *Malat1* knockout models to assess this lncRNA’s role in mediating ethanol-related behaviors. Initial behavioral screening in constitutive knockout mice reported no changes in binge-like ethanol intake or ethanol clearance, and modest female-specific effects on acute functional tolerance. Additionally, female mice significantly reduced daily and cumulative ethanol intake in the EOD-2BC paradigm. A reduction in ethanol consumption in the EOD-2BC drinking assay was also observed following global *Malat1* knockout in adulthood using a tamoxifen-inducible model. These changes in drinking were not attributed to altered taste preference or ethanol clearance/metabolism. Overall, our findings suggest *Malat1* may play a causal, sex-specific role in mediating/modulating alcohol consumption.

## Acknowledgements

We gratefully acknowledge the support of the INIA-Neuroimmune consortium and NIH/NIAAA grants U01 AA020889, R01 AA030257, F31 AA032172, and F31 AA029942.

Figures 1A and 3A were generated under an academic license using BioRender: (Created in BioRender. Gil, D. (2026) https://BioRender.com/86bj754) and (Created in BioRender. Gil, D. (2026) https://BioRender.com/rrb2cto), respectively.

