## Supplemental Tables for "Global deletion of *Malat1* alters alcohol consumption in a sex-specific manner"

**Supplementary Table 1. gRNA target Sequences**

| Name | Sequence |
|---|---|
| *Malat1* TAKO gRNA1 | TCCTCCCAGGAAAACGCAAA AGG |
| *Malat1* TAKO gRNA2 | AAAGGGACACGTCACTCCAC CGG |
| *Malat1* TAKO gRNA3 | AACTTATCTGCGATTTCCTC GGG |
| *Malat1* TAKO gRNA4 | GTTTAGGAGATTGTAAAGGG AGG |
| *Malat1* 5' gRNA | GTTCTCTAGAAATATTCCCG TGG |
| *Malat1* 3' gRNA | GGGTGTAAGGCTTGATTGAG TGG |

Underlined sequence indicates the protospacer adjacent motif

**Supplementary Table 2. PCR Primer Sequences**

| Name | Sequence | Amplicon Size |
|---|---|---|
| *Malat1* Promoter F | CTCCATCTTGTTTCGCA | WT- 933bp |
| *Malat1* Promoter R | AAGTAGGTTAAGTTGACGGCC | |
| *Malat1* qPCR F | GGCGGAATTGCTGGTAGTTT | 197bp |
| *Malat1* qPCR R | AAGGCGTGTACTGCTATGCT | |
| *Malat1* 5' loxP F | TTGGAAAAGACCCACGAAACAA | WT- 375bp<br>loxP 413bp |
| *Malat1* 5' loxP R | ACTTTAGGGGGCGAGGGAAG | |
| *Malat1* 3' loxP F | TGCAATACTGTGTGTAAGTGTGC | WT- 415bp<br>loxP 453 |
| *Malat1* 3' loxP R | CAGGTGAGCAAAATGGTCTCC | |
| *Malat1* ReA F | GGGGAAGACAGTGGGCATTT | Unrecombined: 10.5KB<br>Recombined: 150bp |
| *Malat1* ReA R | TACACCCTGGGCAAAAACATC | |
| *β-Actin* F | GACCTCTATGCCAACACAGT | 150bp |
| *β-Actin* R | AGTACTTGCGCTCAGGAGGA | |
